# Label-free and Multimodal Second Harmonic Generation Light Sheet Microscopy

**DOI:** 10.1101/2020.09.07.284703

**Authors:** Niall Hanrahan, Simon I. R. Lane, Peter Johnson, Konstantinos Bourdakos, Christopher Brereton, Robert A. Ridley, Elizabeth R. Davies, Neveen A. Hosny, Gunnar Spickermann, Robert Forster, Graeme Malcolm, Donna Davies, Mark G. Jones, Sumeet Mahajan

## Abstract

Light sheet microscopy (LSM) has emerged as one of most profound three dimensional (3D) imaging tools in the life sciences over the last decade. However, LSM is currently performed with fluorescence detection on one- or multi-photon excitation. Label-free LSM imaging approaches have been rather limited. Second Harmonic Generation (SHG) imaging is a label-free technique that has enabled detailed investigation of collagenous structures, including its distribution and remodelling in cancers and respiratory tissue, and how these link to disease. SHG is generally regarded as having only forward- and back-scattering components, apparently precluding the orthogonal detection geometry used in Light Sheet Microscopy. In this work we demonstrate SHG imaging on a light sheet microscope (SHG-LSM) using a rotated Airy beam configuration that demonstrates a powerful new approach to direct, without any further processing or deconvolution, 3D imaging of harmonophores such as collagen in biological samples. We provide unambiguous identification of SHG signals on the LSM through its wavelength and polarisation sensitivity. In a multimodal LSM setup we demonstrate that SHG and two-photon signals can be acquired on multiple types of different biological samples. We further show that SHG-LSM is sensitive to changes in collagen synthesis within lung fibroblast 3D cell cultures. This work expands on the existing optical methods available for use with light sheet microscopy, adding a further label-free imaging technique which can be combined with other detection modalities to realise a powerful multi-modal microscope for 3D bioimaging.

## Introduction

In the past decade, light sheet microscopy (LSM) has proven to be a highly effective breakthrough imaging technology and has proliferated very quickly. Its benefits come from the ‘photon-efficiency’ of only illuminating a thin plane that lies in the focus of the detection optics. Out-of-focus excitation is avoided, preventing unnecessary photo-damage of the specimen, and photo-bleaching of fluorophores, in turn allowing higher temporal frequency and/or longer duration imaging. Since little out-of-focus signal is generated, there is inherent z-sectioning^1–4^. The detector captures the full field of view in LSM allowing rapid volumetric imaging when compared to point-scanning systems. Large fields of view are possible with different configurations. Owing to these attributes LSM is found to be desirable for live-cell imaging, long term imaging, and large field-of-view (FOV) imaging of whole small model organisms, such as M. Drosophila^5–7^, C. Elegans^8,9^, and Zebrafish^6,7,10,11^, and for sensitive samples such as embryos^12–14^ and neuronal cultures^15^.

There is, however, a compromise between large FOV and the spatial resolution for a given magnification in LSM. Central to this is the illumination beam profile, which governs the dimensions and properties of the light sheet in 3D space. The default Gaussian profile of a laser beam focused through an objective lens offers high power density at the focal spot, but a very non-uniform illumination along the propagation axis, with a short isotropic region, typically only tens of micrometres. The non-diffracting and self-repairing Bessel- or Airy-type beams^16–20^ maintain near-homogeneous power distribution and resolution over extended lengths in the direction of propagation, leading to increased FOV compared to a Gaussian beam.

A Bessel-type beam can be generated through an axicon lens, or a mask with concentric rings^21^. Self-interference of the wavefronts leads to confinement of the central lobe over long distances, but also generates side lobes that contribute to out-of-focus illumination and reduced resolution. The Airy beam on the other hand can be generated using a tilted cylindrical lens^19,22^ or cubic phase mask^17^. An Airy beam can provide a larger FOV compared to a Bessel beam, whilst also increasing contrast^17^. Both Airy and Bessel beams reduce shadow artefacts in images. With both types of beams, however, deconvolution of the resulting images is needed to recreate diffraction limited images, but this is computationally expensive. Only the relatively uniform central region of an Airy beam is normally used for imaging (**Fig 1a**). It is, however, possible to rotate the Airy beam profile around the axis of propagation to bring the curvature of the beam into the imaging plane of focus^23^. This further extends the field of view because the main lobe remains within the imaging plane despite the beam curvature. Since the light sheet is created by scanning the beam in one plane the curvature is effectively eliminated. The rapid scanning approach of creating a ‘virtual light sheet’ is ideal for multi-photon processes as it provides higher power densities and thus greater signal generation than illumination using a cylindrical lens.

**Figure 1.**
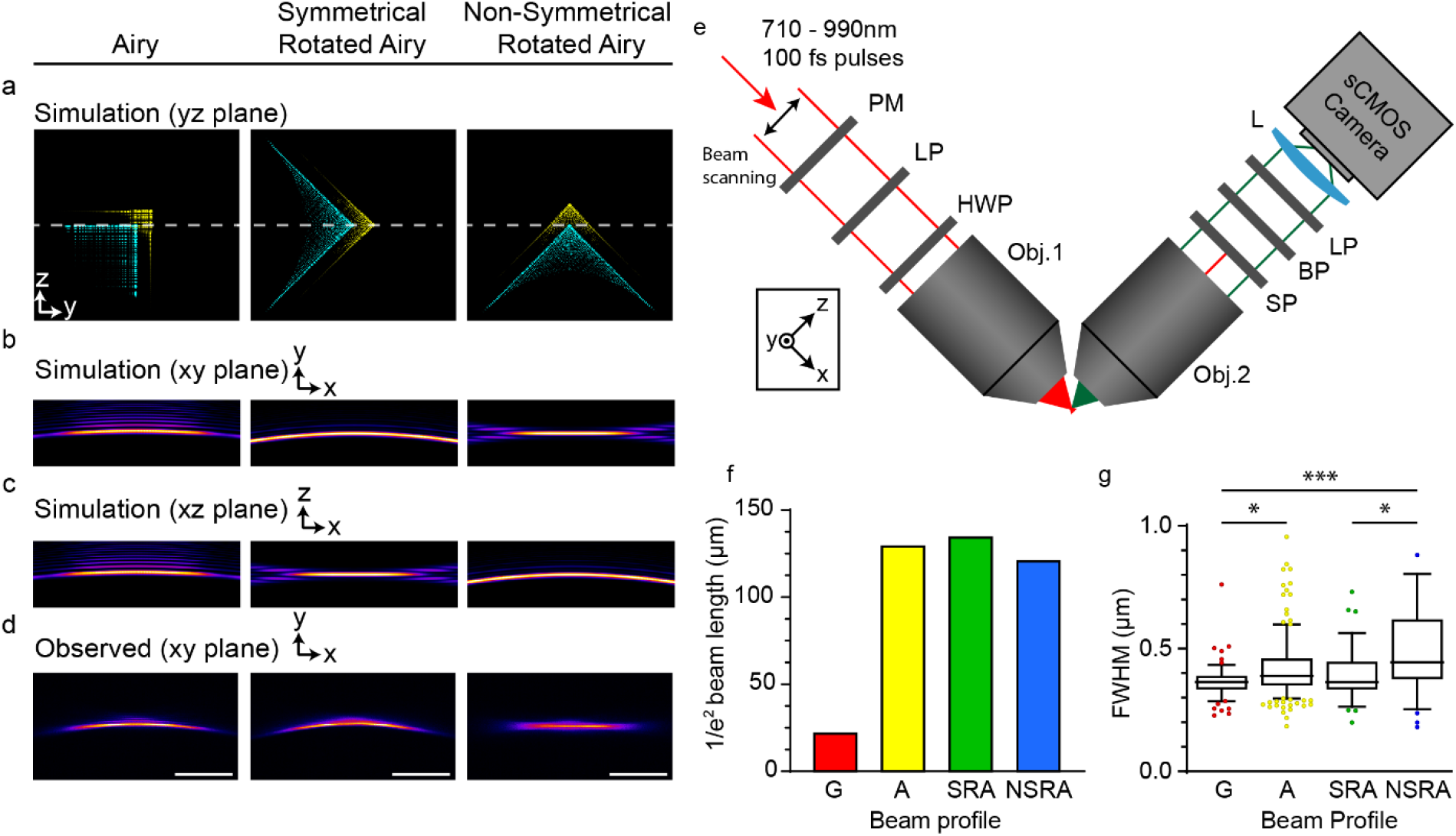
Second Harmonic Generation Light Sheet Microscope (SHG-LSM) (a) Simulated beam profile for the Airy, Symmetrical Rotated Airy (SRA) and Non-Symmetrical Rotated Airy (NSRA) beams in the y-z plane (Cyan, x=0μm; Yellow, x=±100μm). Dashed lines indicate focus of the detection objective. (b,c) Simulated beam profiles for two photon fluorescence in the x-y (b) and x-z (c) planes. (d) Observed beam profile in the x-y plane using two-photon excitation of 100μM FITC. Scale bar represents 30μm. (e) Illustrative imaging path of Second Harmonic Generation Light Sheet Microscope (SHG-LSM); PM, Cubic Phase Mask; LP, Linear Polariser; HWP, Half-wave plate; SP, Short-pass filter; BP, Band-pass filter; Inset shows the global coordinate system used throughout this work. (f) Measured beam 1/e^2^ beam profile of the beams in ‘e’, as well for a Gaussian beam (G, no Airy phase mask). (g) Experimental lateral FWHM of 60nm particles imaged using the beams in ‘f’. (See Supplementary Table 1 for full experimental details)

Multi-photon light sheet microscopy typically involves excitation of fluorophores using a near-infrared (NIR) pulsed laser and emission is at visible wavelegnths^7,16,24^. The NIR excitation allows for improved penetration into biological samples and minimises scattering compared to visible excitation^25^. Two-photon light sheet fluorescence microscopy (2P-LSFM) has been used for live imaging of zebrafish development^7^, and more recently three-photon light sheet fluorescence microscopy (3P-LSFM) has been used in conjunction with Bessel beam illumination for 3D imaging with high-contrast and low photodamage in highly scattering cell spheroids over a large FOV^26^. While with one-photon (1P) excitation the out-of-focus side-lobes in Airy and Bessel beams cause a reduction of image contrast, with 2P excitation, due to the quadratic dependence on incident intensity, this issue gets largely resolved.

Second Harmonic Generation (SHG) microscopy is also a multi-photon imaging method, which is finding increased application in biomedicine. SHG is a parametric nonlinear optical process occurring in non-centrosymmetric structures where two photons get combined resulting in a photon at twice the frequency of the input photons (ω ⟶ 2ω)^27^. SHG is not prone to photo-bleaching or photo-damage that affects fluorescence-based techniques as it involves non-resonant electronic transitions^28–30^. Warious dyes^29–31^ and biological structures such as fibrillar collagen^32^, myosin^33^ and microtubules^34^ are highly SHG active. Thus, collagen fibres can be imaged without labelling, which finds application in cancer scoring and collagen type/orientation identification^35–38^. The orientation of collagen fibres can also be determined since SHG signals are dependent on the polarisation of the excitation^39^.

SHG is a coherent process, hence, apart from being dependent on the polarisation state of the incident beam the signals are also highly directional^40^. SHG signals propagate largely in the forwards and backwards direction, which has been established theoretically^41^ and experimentally^42^. In biological samples, however, SHG emission directionality depends on a number of factors, including the materials properties, number of scatterers, scatterer spacing, size and orientation of scatterers in the focal field, as well as the polarisation state of the incident beam and excitation intensity^43^. For collagen fibrils their orientation relative to polarisation axis of excitation affects SHG emission directionality^44^. Whilst SHG signal intensity does not depend on the numerical aperture (NA) of the illumination objective^29,31^, the NA does affect the directionality; for NA<0.8 the angle (θ_*peak*_) of maximum SHG signal scales with the illumination angle (θ_*NA*_) determined by the NA as given below by Equation 1^45^:

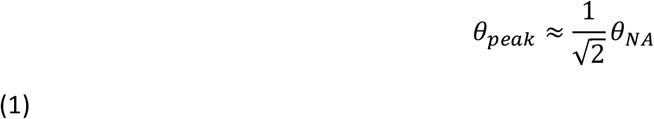

This means that for a single scatterer, the lobes of SHG signals would be expected at a proportionally shallower angle than determined by the NA of the illumination objective. For SHG signal generation the phase matching condition (Equation 2) within a material must also be satisfied:

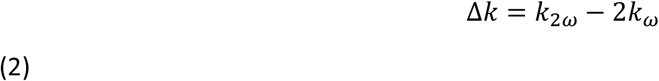

Where *k*_2ω_ and *k*_ω_ are the wave vectors of the SHG and incident photons^36^. In the case of perfect phase matching, i.e. where phase mismatch Δ*k* = 0, emission propagates in the forward-scattered and back-scattered (F_SHG_, B_SHG_) directions^32,36,46^. Second harmonic signals, however, can be generated in other directions In the case of imperfect phase matching, where Δ*k* ≠ 0 such as in biological tissues^43^. Importantly, however, SHG signals are directional rather than isotropic, like in fluorescence emission (1PF or 2PF). Existing 3D imaging approaches use a variety of mechanical or computational methods^39,46–48^, but all generate data through point-scanning, and use only a forward- and/or back-scattered detection geometry for SHG signal collection. Since LSM uses an orthogonal signal collection configuration instead of collinear excitation and detection, it presents a frontier for SHG imaging. Provision of SHG on a LSM however, is much desirable as a modality to rapidly image the 3D distribution and organisation of harmonophores such as collagen fibrils within tissues, which amongst other things has implications for cancer screening^37^.

In this study we demonstrate SHG imaging on a light sheet microscope. We use a rotated Airy beam for rapid deconvolution-free volumetric SHG imaging (300 μm ×150 μm × 80 μm) providing large FOV without compromising on spatial resolution as compared to Gaussian and conventional Airy beams. We find that there is an improvement in the field of view and resolution at over >100 μm distances. The SHG signals are duly verified using their wavelength and polarisation dependence. We show that SHG imaging in the typical orthogonal detection configuration of an LSM allows imaging of multiple sample types including collagenous tissue, SHG-active dye intercalated to cell membranes, and from within 3D cell spheroids. Additional information can be obtained by using the polarisation dependence of SHG signals on the LSM. 2PF signals can also be acquired providing multimodality combining both unlabelled and labelled samples. To demonstrate the utility of the multimodal SHG-LSM for biomedical studies we carry out a study of 3D lung fibroblast spheroids, confirming clear changes in the SHG signal when collagen production is promoted. Given that structure readouts of collagen provided by SHG imaging can allow diagnostic and prognostic information in cancer^49^ and can also be markers of fibrotic lung disease^50^ our 3D SHG-LSM approach can be transformative both for medical research and drug screening.

## Results

To realise SHG imaging on a light sheet microscope we developed a multi-photon system that utilised a cubic Airy phase mask (APM) in Fourier space to generate an Airy beam at the sample (**Fig 1a**). A pulsed NIR laser beam is scanned laterally in the y-axis by a resonant scanning mirror to give a time-averaged light sheet in the imaging plane of the detection objective.

### Symmetrical Rotated Airy improves resolution across FOV

We first tested the effect of rotation of the Airy beam profile on the field of view (FOV) and the spatial resolution. We achieved this in our setup by rotation of the Airy phase mask (APM, **Fig 1a**). The Airy beam profile consists of a main lobe, and then a series of secondary lobes of diminishing power. We estimated that the depth of field of our detection objective limited signal collection to <1 μm of the focus, whilst the distance from the primary lobe to the nearest secondary lobe was <2 μm (**Fig S1**). In addition, we are exciting multi-photon processes (SHG, TPF) that have quadratic power dependence for signal generation. Hence, the contributions from more distant secondary lobes can be ignored as they are significantly weaker (for two-photon excitation the secondary lobe contains ^~^20% of the intensity compared to the main lobe). Furthermore, since the secondary lobe lies outside of the depth of field of the collection objective, out-of-focus contributions to signal are reduced even further. In the standard Airy beam profile one set of side lobes lie in the imaging plane, thus contributing to in-focus illumination as the beam is scanned in the y-axis^17^. However, the secondary lobes contribute to background signal away from the imaging plane, reducing image contrast (**Fig 1b**, Airy). Depending on the Airy beam properties, the lobe structure can be used for recovery of image contrast by deconvolution with the PSF^17^. However, if the lobe structure is such that contributions from side lobes is sufficiently small, deconvolution is not necessary.

For developing a deconvolution-free SHG-LSM method we tested two alternate orientations: with the secondary lobes placed symmetrically above and below the imaging plane, termed Symmetrical Rotated Airy (**Fig 1b**; SRA), or with all secondary lobes being either above or below the imaging plane, termed Non-Symmetric Rotated Airy (**Fig 1b**, NSRA). A recent publication by Hosny et al. uses the term Planar Airy Light Sheet which is the same case as our Symmetrical Rotated Airy^51^. In this work, we distinguish the two beam types by their symmetry about the illumination plane. The curvature of the beam causes a shift in the position of the central lobe as it propagates along the z-axis, with only the SRA profile maintaining the main lobe in the focus across its entire field of view. This was confirmed by both simulations (**Fig 1 b-d**) and experimental measurements (**Fig 1e**). Analysis of the empirical data showed that the FOV was extended slightly (4%) using the SRA profile, whilst it was reduced slightly (7%) in the NSRA profile, when compared to the Airy profile (**Fig 1f**; Airy, 130 μm; SRA 136 μm; NSRA 121 μm). Larger improvements to effective FOV are expected when using different combinations of objective and Airy beam profile, with an improvement of up to 33% demonstrated^52^.

To determine the effect that these profiles have on resolution we imaged fluorescent beads (Fluoresbrite^®^ YG Microspheres 0.10μm, Polysciences Inc.) and calculated the full-width at half-maximum (FWHM) at different positions along the x-axis. We found that the Gaussian beam achieved a FWHM of 380 ± 32 nm in the centre of the FOV (x=0μm), and a significant difference was measured with the Airy (p > 0.05) and NSRA (p > 0.001) profiles, of around 400 nm and 450 nm respectively. The FWHM for the SRA beam was not significantly different from the Gaussian beam, at around 380 ± 89 nm (**Fig 1g**). The resolution measured in the centre and at the edges of the FOV were not significantly different in all cases, apart from the NSRA beam profile (**Fig S2**). The out-of-plane curvature of the NSRA beam away from the beam centre is the likely cause of this. Results of simulations for each beam type are presented in Figure S1. The almost uniform MTF for the SRA beam demonstrates the spatial resolution should be invariant across the 150 μm FOV in the range specified, whereas the MTF for the NSRA beam shows a significant reduction in achievable resolution away from the centre of focus. The SRA beam therefore yields an increase in effective field of view, without compromising on resolution, and so was used in the remainder of this study.

### Second Harmonic Generation Light sheet Microscopy (SHG-LSM)

SHG signal propagation is predominantly in the forwards and backwards direction and hence adapting it for an orthogonal detection configuration in a light sheet microscope is counter-intuitive. For single particles, or ideal SHG scattering materials e.g. crystals excited by linearly polarised light the far field SHG signal is conical in shape and comprises of two opposing lobes. This cone angle is a fraction of the NA of the illumination objective, and thus does not fall within the collection angle of the detection objective. However, for non-uniform scatterers, such as biological specimens, the phase-matching condition is relaxed and side scattering of SHG is expected to be significant^45^. Hence, some signal can be captured especially with a high NA objective to make SHG-LSM possible (**Fig 2a**).

**Figure 2.**
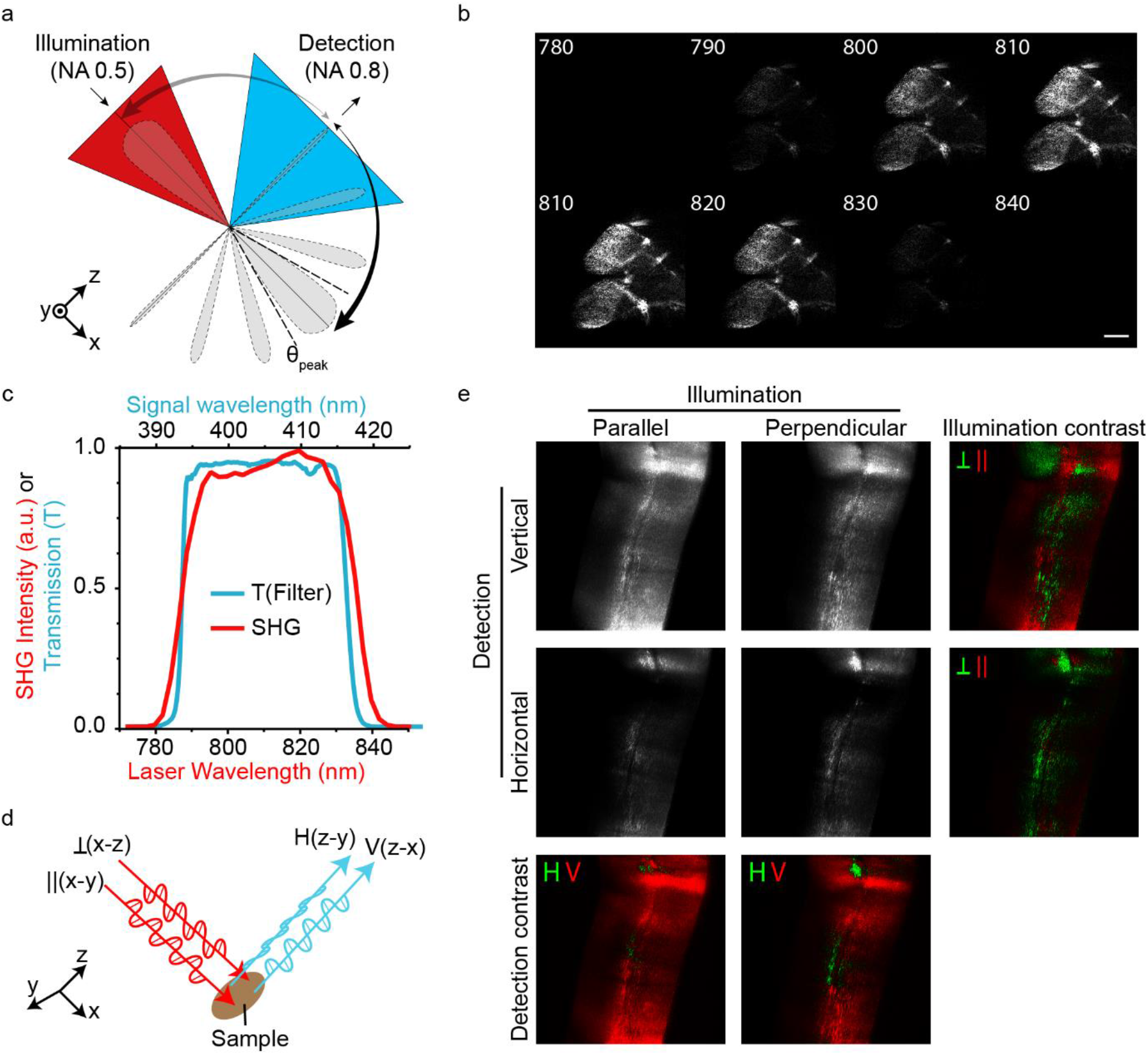
Wavelength and polarisation dependence of LS-SHG signal. (a) Schematic showing the relative illumination (red) and detection (blue) angles for our system. θ_peak_ contains the predicted maximal signal intensity for perfect scattering samples. Non-perfect scattering materials may produce weaker scattering at larger angles. (b) Images from a wavelength scan of rat-tail collagen, showing 10nm increments, with detection through a 405±10nm filter. (c) Plot of data obtained from ‘b’ at 2nm increments, overlaid with the transmission profile of the 405±10nm filter. (d) Schematic depicting the polarisations of light used in the illumination and detection paths. Perpendicular (⊥) and parallel (∥) refer to the orientation of the fast axis of polarisation relative to the plane of the light sheet. Horizontal (H) and vertical (V) refer to the two orthogonal detection filters applied to the detection path (e) SHG signal from rat-tail collagen imaged using the illumination and detection polarisation states shown in ‘d’. Detection contrast shows the difference between two channels, where positive values are in red and negative values are in green. (See Supplementary Table 1 for full experimental details)

In order to verify that SHG imaging is possible on an LSM we used a section of fixed rat-tail tendon, rich in type-I collagen that is highly SHG active^36^. We obtained images with excitation wavelengths in the range 730 nm to 860 nm with detection through a 405 ± 10n m bandpass filter. We observed that the same images were generated only between 790 nm and 830 nm (**Fig 2b,c**), as these wavelengths corresponded to frequency doubled signals through the bandpass filter. No signal was detected outside of this range. The slight broadening of the signal profile relative to the bandpass filter can be explained by the spectral bandwidth (Δλ) of the 100 fs pulsed laser as given by Equation 3 below:

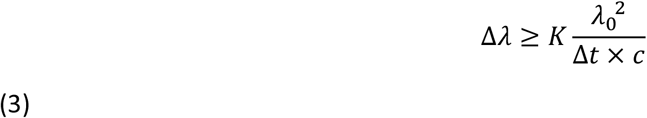

Where Δ*t* is the temporal pulse width, λ_0_ is the central wavelength of the pulse, *c* is the speed of light, and K is a constant describing the time-bandwidth product for a Gaussian pulse shape (*K* = 0.441 for a Gaussian pulse). Δλ is calculated to be 9.4 nm. Since SHG is frequency doubled, we therefore anticipate a spectral bandwidth of 4.7 nm in the SHG signal. Consistent with this, there was a gradual change in signal from collagen over ^~^5 nm at the bandpass filter cut-on and cut-off wavelengths. While this experiment indicated that the signal to be the result of second harmonic generation, we wanted to verify further as SHG will be highly polarisation sensitive unlike cellular auto-fluorescence.

### Polarisation Dependence of Orthogonal SHG signal

SHG generation requires that the incident light polarisation is aligned with the SHG-active structures in the sample, and the SHG signal generated is also polarised. With respect to the image plane that is detected (the x-y plane), we used two illumination polarisations, Parallel (electric field oscillation in the x-y plane), or Perpendicular (electric field along the z-axis). Similarly detection polarisation was defined as being Vertical (x-z plane), or Horizontal (y-z plane) as depicted in **Fig 2d**. These polarisations were achieved using a linear polariser coupled with a half-wave plate in the illumination path, and linear polarising filters in the detection path (**Fig 1a**). We then probed a sample of rat tail collagen using each permutation of the above polarisations. We found that different regions responded to particular combinations (**Fig 2e**), presumably reflecting the predominant underlying collagen fibre orientation in that region. This was probed further by varying input polarisation through its full range and measuring the signal from the anisotropic sample (**Fig S3**). No significant difference was measured in SHG signal from collagen using left- and right-handed circular polarised illumination (**Fig S4**). Collagen fibril orientation varies within the illumination plane and relative to the illumination plane. As polarisation of the input light is rotated about the illumination axis, the typical dumbbell response is observed. The major axis of this dumbbell changes with collagen fibril orientation, as does the intensity of the response. Together the wavelength- and polarisation-sensitivity of the signal indicates it is the SHG signal that is collected orthogonally from the light sheet microscope.

### Compatibility of SHG-LSM for Multi-Modal Bioimaging

We next performed SHG on live cells, using mouse oocytes as a specimen. These large (80 μm diameter) and spherical cells were loaded with the SHG active dye FM4-64, which intercalates into plasma membranes, thus forming an annular-sphere of the SHG dye around the cell. The FM4-46 molecules therefore have a predictable directionality, being orientated normal to the membrane surface at any given point. Wavelength-scan imaging of an oocyte revealed that the SHG signal could be visualised at the anticipated wavelengths (790-830 nm, **Fig 2c**) and in the expected region at the periphery of the cell (**Fig 3a**, 790 nm to 830 nm, inset shows the one-photon excited fluorescence signal from FM4-64 in the same cell). The strongest signal was from autofluorescence of NAD(P)H in the cytoplasm, however we found that contrast was possible for SHG at wavelengths from 810 nm to 830 nm, where NAD(P)H excitation is minimal, allowing the two to be easily separated (**Fig 3b**). Further to this we could also simultaneously image the DNA intercalating dye Hoechst, which has two-photon excitation within the same wavelength range. We utilised a K-means clustering technique to segment the image data by grouping pixels with similar spectral profiles together. This resulted in clear separation of the SHG signal, and the two fluorescent signals in live cells (**Fig 3c**), showing that light-sheet SHG imaging is compatible with other labelled, or label-free imaging modes.

**Figure 3.**
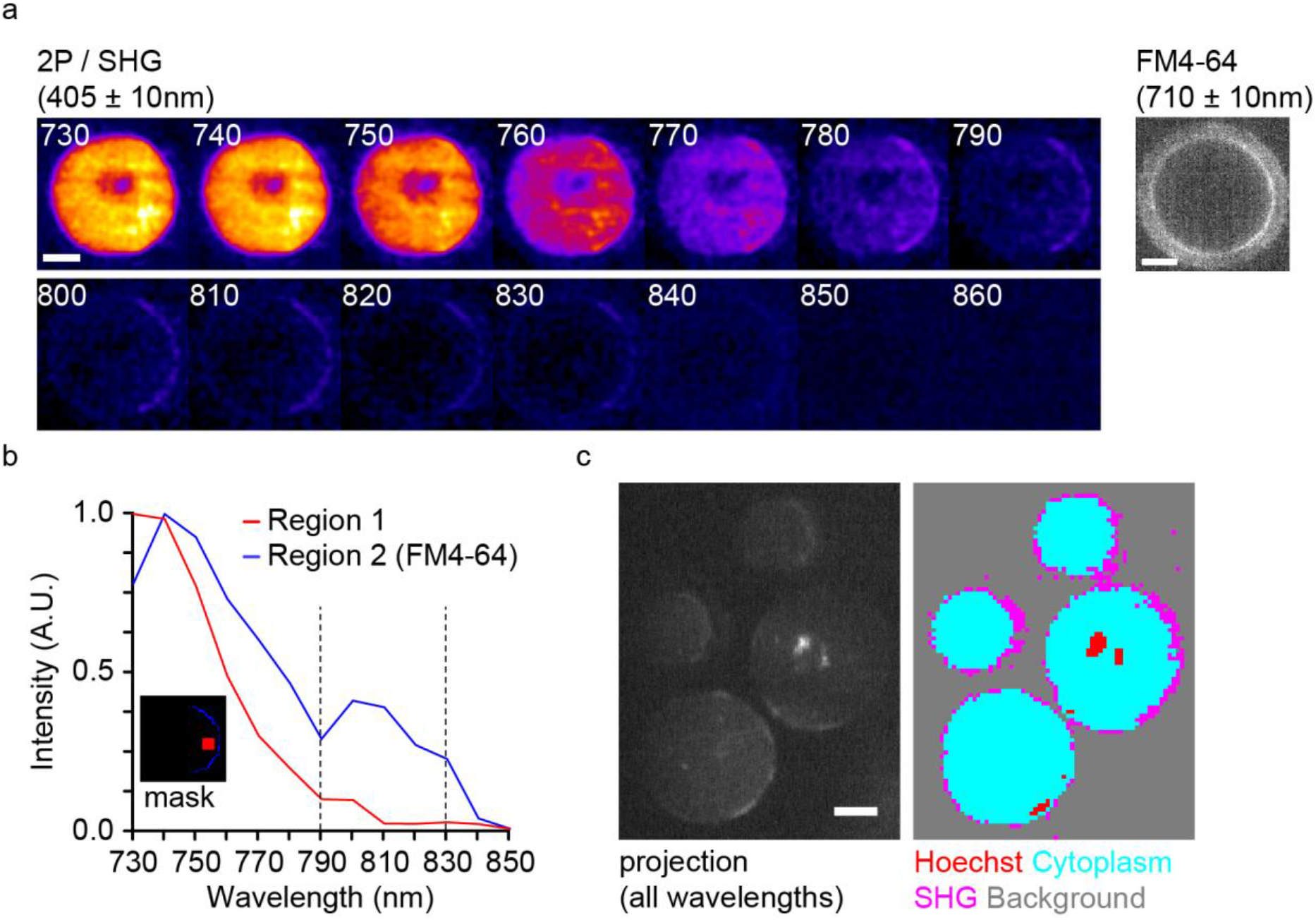
Spectral and spatial separation of 2P-excited autofluorescence and SHG signals. (a) Emissions through 405 ±10 nm bandpass filter with fs-pulsed laser excitation in range 730-860 nm from oocytes treated with FM4-64 (10μM). Greyscale image shows 1PF signal from plasma membrane FM4-64 dye (exc: 488nm, Em:710±10nm). (b) Normalised excitation spectrum from cytoplasm and from plasma membrane using the regions depicted in the insert. Dashed vertical lines show the expected SHG emission range for the detection filter (405±10nm). (c) K-means clustering used for separation of spectrally distinct image regions. (See Supplementary Table 1 for full experimental details).

We next wanted to take advantage of the unique spherical plasma membrane of the oocyte, which provides us with all possible orientations of the SHG dye in one sample. We acquired 3D image stacks through the oocytes loaded with the dye, recorded the one-photon fluorescence signal of the dye as well as the SHG signal (**Fig 4a**). Using perpendicularly polarised illumination we were able to visualise two regions of the oocyte plasma membrane that generated SHG signal. Projection of the images in the x-z plane revealed that the two regions were opposite each other, at 45° and −135° to the incident light direction in the xz plane (**Fig 4b, c**). We reasoned that at only these specific orientations were two criteria for light sheet SHG fulfilled: firstly, that SHG scattering from the dye molecules was possible at that orientation, and secondly that the dye orientation caused the SHG emission to fall within the detection cone of the objective (**Fig 4d**). When we changed the illumination polarisation by 90°, such that it was parallel to the illumination plane, the SHG signal was lost at these locations (**Fig 4e**), and in addition, did not show up at any other location on the oocyte plasma membrane (data not shown), suggesting that in this case one of the two criteria was not fulfilled.

**Figure 4.**
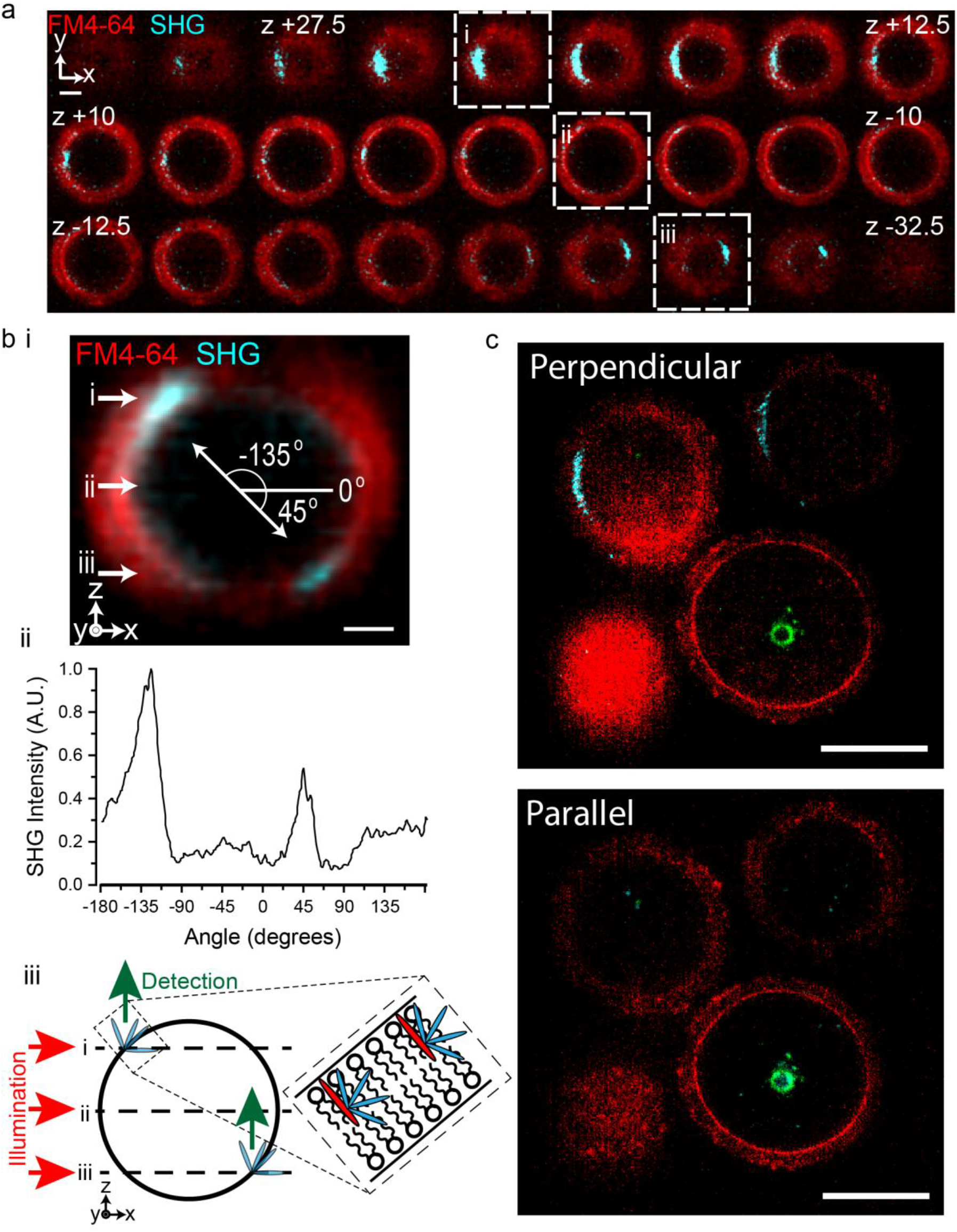
Lightsheet SHG signal depends on harmonophore orientation and input polarisation state. (a) 3D image stack of an oocyte stained with FM4-64, showing 1PF (red; ex:488nm, em:710±10nm) and SHG (cyan; illumination:810nm, em:405±10nm). Three z-positions, symmetrical about the oocyte centre, are highlighted (i,ii,iii). (b) (i) Projection of the image stack from ‘a’ in the y axis, showing the location of the three indicated z slices (i,ii,iii). The direction of illumination propagation (x axis) was used to define 0 degrees. (ii) SHG intensity at the cell membrane for all angles from the centre of the oocyte, using the coordinate system defined in ‘b’. (iii) Schematic showing the dependence of SHG signal collection on the relative angle of the emitter. (c) A single plane from an imaged volume of an FM4-64 stained oocyte with either perpendicular (left) or parallel (right) polarisation of illumination (red, 1PF; cyan, SHG). Scale bars represent 20 μm (a, b(i)) and 50 μm (c). (See Supplementary Table 1 for full experimental details)

### SHG-LSM imaging of 3D Tissue-Engineered Model of Human Lung Fibrosis

Most multicellular organisms require the use of an extracellular matrix (ECM) to grow and define their 3D shape. ECM provides structural as well as biochemical support to the cells they enclose, defining directionality within tissues as well as the boundaries between tissues^53^. Its study is therefore essential to understand a wide range of biological and medical problems, e.g. tissue invasion by cancer cells, or bone morphogenesis. ECM comprises an aggregated mesh of proteins and glycoproteins, with the most abundant member being collagens^36^.

To demonstrate that SHG-LSM is well-suited for large-scale 3D imaging and that the technique can be used to detect changes in collagen we carried out experiments using our long term 3D tissue-engineered model of human lung fibrosis^50^, which forms 3D spheroids using human lung fibroblasts and produces structured incorporated ECM including cross-linked fibrillar collagens. We performed label-free 3D imaging of the cells and the ECM within the 3D spheroids (diameter 450-800 μm), targeting the cellular two-photon autofluorescence and SHG signals, respectively (**Fig 5a**). The autofluorescence signal included strong localised luminescence from Nanoshuttle particles (used for manipulation of spheroids). The SHG signals imaged fibrillar collagen in the ECM. Thus multimodal 3D imaging with 2PF and SHG allowed us to image fibroblasts and ECM respectively (**Fig 5b**). In this label-free multimodal SHG image acquisition we could image cells and ECM in the spheroids across a large effective total volume at high spatial and temporal resolution (see Methods). Our current setup has a single detector but with filtering it is possible to acquire both 2PF and SHG signals simultaneously.

**Figure 5.**
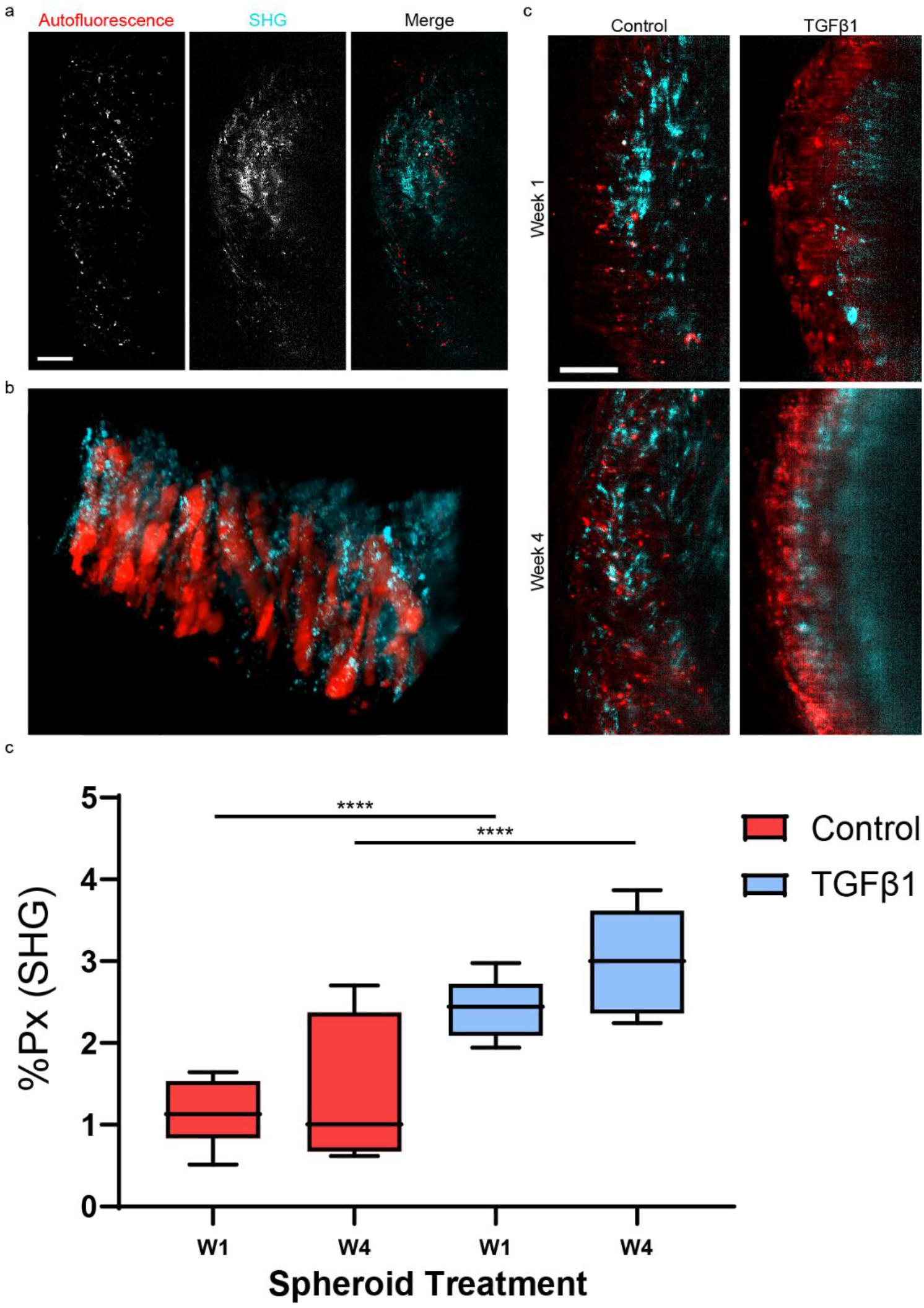
SHG-LSM for multiphoton imaging of lung fibroblast spheroids. (a) Composite images of a lung fibroblast spheroid after 1 week of culture, showing autofluorescence (cyan) and SHG (red) signals. Each image is a single frame comprised of 2 volumes stitched in the y-axis. Scale bar represents 50μm. (b) 3D rendering of an unlabelled lung fibroblast spheroid (autofluorescence, red) and showing extracellular matrix (SHG, cyan). See also Supplementary movie 1 (c) development of lung fibroblast spheroids with or without TGFβ-1 addition to the culture medium (autofluorescence, red; SHG, cyan). Scale bar represents 50 μm. (d) The SHG signal from the ECM was compared across two different treatments and two timepoints. One-way ANOVA on SHG signal obtained from spheroids shows a significant difference between control and TGFβ1 treatments (P < 0.0001) for both week 1 and week 4 timepoints. (See Supplementary Table 1 for full experimental details)

We further used SHG-LSM to assess temporal changes in collagen content of the 3D spheroids following culture for 1 or 4 weeks in the presence or absence of the pro-fibrogenic cytokine TGFβ1 (3 ng mL^−1^). All spheroids were imaged under the same conditions, to ensure differences in SHG signal were attributable to changes in collagen alignment, amount, density or structure. Image stacks were acquired in 3D (see **Fig. 5b**) and in both 2PF and SHG channels at two different positions within each spheroid. Low average illumination power (<100 mW) was used to prevent damage to the spheroids. Consistent with the known induction of collagen by TGFβ1^54^, significant differences in collagen SHG signals relative to controls were observed for spheroids treated with TGFβ1 (P>0.0001) at both week 1 and week 4 (**Fig. 5c**).

## Discussion

In this work we have demonstrated SHG imaging on a light sheet microscope in a systematic manner and with multiple types of biological samples. We verified unambiguously that the signals generated are due to second harmonic generation. Whilst theory dictates that this will be an inefficient process, we believe this is somewhat offset by the inherent ‘photon-efficiency’ of the light sheet when compared to point-scanning systems. Further to this, the increase in speed and volume that is possible when combined with the light sheet configuration greatly adds to its usefulness.

Here we have demonstrated that SHG imaging over large volumes (0.027mm^3^) can be performed rapidly (e.g. 0.081 mm^3^ min^−1^ in our current system). We further tested various Airy beam configurations for SHG-LSM. We have also demonstrated that rotation to a standard Airy beam profile gives a modest increase in the useable field of view, with no loss of resolution. Creation of an Airy beam profile in a light sheet microscope is straightforward to implement, either by the positioning and rotation of a cubic phase mask^17,52^ or cylindrical lens^19,22^, or by design of SLM pattern^55^. Experimentalists using an Airy beam light sheet microscope should strongly consider use of the symmetric rotated Airy beam (SRA) beam profile rather than the standard Airy beam profile to realise the increase in useable FOV and resolution uniformity afforded by this beam type. The SRA beam profile provides an increase in useable FOV compared to the standard Airy beam and NSRA, as the main lobe remains in the illumination plane across the FOV as shown in **Fig. 1a**.

The use of a Symmetrical Rotated Airy beam allows for deconvolution-free direct imaging of SHG-active structures across a large FOV, while still providing high resolution comparable to a Gaussian beam. Critically, this high resolution is maintained across the full width of the FOV, which can be (with an appropriate choice of Airy beam α-value) >200 μm in the x-axis. Towards the edges of the FOV, the focus of the main lobe for the Airy and non-Symmetrical Rotated Airy (NSRA) beams is found outside of the imaging plane, causing an appreciable reduction in resolution away from the beam centre. The modulation transfer function (MTF) of the objective system was simulated for all beam types, (**Fig S1**) indicating both that the achievable resolution in x is ^~^440 nm, and that the resolution from the SRA beam is more uniform across the FOV than that of the NSRA and Airy beam profiles. At the centre of the FOV (x = 0 μm), the achievable lateral resolution of the SRA beam is better than 400 nm, identical to both the Airy and NSRA beams, but over a wider range. Dholakia and co-workers used 1P fluorescence with a standard Airy beam illumination profile on an LSM and obtained an axial resolution of 1.9 μm and a lateral resolution of 1.5 μm across a 200 μm FOV^17^. Further work performed by Hosny and co-workers used a Symmetrical Rotated Airy beam for illumination of the FOV with 2PF detection, achieving 0.83 μm lateral and 3.69 μm axial resolution for the SRA beam, and 0.91 μm lateral and 3.74 μm axial resolution for the Airy beam. This resolution was achieved across an effective FOV of 415 μm which, compared to the FOV of the standard Airy beam profile of 311 μm, represents an improvement of 33%.

With the SRA we showed that SHG imaging with orthogonal collection in an LSM is possible. The wavelength and polarisation dependence confirmed unambiguously that the signal was due to SHG. Alignment of SHG harmonophore with excitation electric field determines both the strength and the direction of SHG emission. The spherical oocyte cells allowed us to probe SHG excitation and emission on the LSM. SHG emission direction changes depending on harmonophore alignment relative to the excitation electric field, and SHG signal is observed when there is alignment of the orientation of a harmonophore with the polarisation state of illumination^56,57^. The FM4-64 dye molecules are orientated parallel to the plasma membrane at every position on the oocyte surface, covering all possible orientations of the dye molecule, and thus are ideal for investigating the orientation dependence of the SHG signal from a simple harmonophore^58,59^. The signal dependence on input polarisation state agrees with Malkinson and co-workers on SHG-SPIM, who found that nanocrystal orientation and illumination polarisation both affect the orthogonally-detected SHG signal distributions measured from randomly orientated KTP and BaTiO_3_ nanocrystals suspended in agarose^57^. Average SHG signals from these nanocrystals were significantly reduced when the polarisation axis was parallel to the illumination plane^57^, in agreement with the FM4-64 dye results presented in this work. It is most likely that SHG is still generated with the parallel polarised illumination but is outside the detection volume of our LSM configuration.

The use of image stitching extends the volume arbitrarily in the y-axis. The resulting images are suitable for quantitative studies, in our case showing changes in extracellular matrix production during the growth of lung fibroblast spheroids in culture or showing collagen directionality in rat tail samples. We acknowledge that due to the dependence of SHG on sample-orientation, that emission from some SHG active structures will not be captured orthogonally. This could be addressed in future by sample rotation within the light sheet, as is common in many SPIM setups.

Aside from collagen detection, SHG imaging may be used for imaging the mitotic spindle (microtubule based structures present only in mitosis), e.g. for assessment of mitotic stage in cultured cells, or for spindle positioning in IVF, cancer tissue differentiation, and bone sample composition.

Since SHG emission, unlike fluorescence, is a scattering process, it is not confined to any particular excitation and detection wavelengths. Therefore, it is possible to adjust the illumination wavelength, and collection bandpass window to fit around other experimental requirements, e.g. to avoid overlap with autofluorescence or other sources of emission. The SHG scattering spectrum is also very narrow, being defined by half of the illumination wavelength bandwidth, meaning exclusion of other signals can be easily achieved with a narrow bandpass filter. SHG-LSM is therefore likely to fit well with other imaging modalities, and indeed, adding a dichroic mirror to the detection path allows simultaneous collection of SHG signal along with fluorescence, further reducing sample light exposure. SHG can be added to existing light sheet microscopes that have a pulsed NIR laser by adding an appropriate bandpass filter to the detection path, and ideally adding simple polarisation control optics (half-wave plate) to the illumination path.

In this work we use an SRA beam to provide an extended high-resolution FOV for sample illumination, and demonstrate the first use of SHG-LSM for label-free imaging of native contrast from biological structures (collagen, ECM). Generation of contrast without the need for exogenous contrast agents (e.g. dyes, fluorescent proteins), termed label-free imaging, is likely to become more prominent in biological and medical research as the desire to understand system behaviour in live cell models, whilst minimising perturbation or behaviour/function changing interventions (such as labelling), increases. In addition, advances in culture systems to better represent in-vivo conditions push scientists towards 3D culture systems, or multi-tissue organoid cultures, which require larger imaging volumes and greater depth penetration. Taken together, we believe that label-free imaging by light sheet microscopy, with an emphasis on the use of multi-photon and near-infrared excitation for imaging, will be a valuable future research tool as it can accommodate the growing need for rapid volumetric imaging, label-free and low photo-toxic imaging modalities. Our demonstration of SHG-LSM is a vital step in this direction to achieve multimodal label-free imaging on a light sheet microscope.

## Materials and Methods

### Multi-Photon Light Sheet Microscope

For this investigation we used an Aurora Airy Light Sheet Microscope (M-Squared Life) as the platform system. The laser output from a fs-pulsed Ti:Sapph laser (MaiTai-BB, 710-990 nm, 80MHz) was coupled into the entry port of the microscope. An Airy beam is created and incident on the sample through the illumination objective (Olympus UMPFLN20XW, 20x, 0.5 NA, WD 3.5 mm). The light sheet is created by laterally scanning this beam in the imaging plane of the detection objective (Olympus LUMPLFLN40XW, 40x, 0.8 NA, WD 3.3 mm). Typically, full-FOV images were acquired across 300 μm in z with a 2 μm slice spacing, and within 10 min per modality (2PF, SHG, etc.).

In the SHG imaging mode, the laser wavelength was set to 800 or 810 nm, and a bandpass filter centred at ^~^400, or 405 nm was used. In the 2PF imaging mode, the laser wavelength was set to 740 nm, and a 520±20 nm bandpass filter was used.

Polarisation control in the illumination path was achieved using a half-wave plate allowing continuous control of the incident polarisation angle. For circular polarisation, a quarter-wave plate was placed in the incident beam path with the fast axis placed at −45° or +45° for left- or right-handed circular polarisation respectively.

Samples were positioned in the camera field of view using a 3-axis linear translation stage, and 3D image stacks were acquired using a motorised linear stage orientated along the xz axis (perpendicular to the imaging plane).

### Sample Preparation

All experiments involving animals were carried out in accordance with the Animals (Scientific Procedures) Act 1986 set out by the UK Home Office as well as all local regulations.

Rat tail tendon was dissected and immediately fixed with 1% paraformaldehyde in PBS for 1 hour. Collagen-containing fibres were removed from the rat tail and stored in 4% paraformaldehyde until needed for imaging. Before imaging, fibres were washed with DI water, and cut into ^~^3 mm pieces using a scalpel.

For experiments involving oocytes, the cells were harvested as described previously^60^, briefly 3-4 week old MF1 mice were hormonally primed (10IU PMSG, Cenataur Services) to increase oocyte yield, and GV stage oocytes were collected 48 hours later by dissection of the mouse, and liberation of the oocytes from the ovaries into M2 media^61^ under a dissection microscope. Oocytes were stored in the dark, in M2 media under paraffin oil on a 37°C heat block until needed.

### Lung Fibroblast Spheroids

Human lung fibroblast spheroids (approximately 500 - 800 μm diameter) were cultured according to our previously reported 3D in vitro spheroid model of lung fibrosis methodology^50,62^ which enables the study of all aspects of collagen supra-molecular assembly, with the methodology adapted to incorporate NanoShuttle (Greiner Bio-One, Abingdon, UK) technology according to manufacturer’s instructions. Briefly, primary human lung fibroblasts were established under the approval of the Southampton and South West Hampshire and the Mid and South Buckinghamshire Local Research Ethics Committees (ref 07/H0607/73) from macroscopically normal lung parenchyma tissue of patients undergoing early stage lung cancer resections. The lung fibroblasts were grown to 80% confluence and then labelled overnight with NanoShuttle-PL which contains gold, iron oxide and poly-L-lysine to magnetize cells to promote uniform spheroid formation. The following day cells were seeded in a 96 well Greiner-Bio-One cell-repellent surface plate. After putting the plate on top of the Greiner-Bio-One magnetic drive for 1 h, followed by incubation for 24h to allow cells to form 3D spheroids, cells were changed to long term DMEM/F12 media^50^ in the presence or absence of 3 ng/ml TGF-β1 (R&D Systems, Abingdon, UK). Media was replenished three times per week, and cell spheroids were harvested at 1 and 4 weeks and fixed in 4% paraformaldehyde before imaging.

### Sample Mounting

Custom sample holders were prepared from a microscope slide, a small cylinder of PDMS (d = 5 mm, h = 5 mm), and a small weighing dish with 5 mm central hole. The dish was attached to the microscope slide using double-sided tape, and the PDMS cylinder attached directly to the tape through the hole. Low-melting point (LMP) agarose (1% w/v, Sigma, A9414) was prepared in DI water, and left to cool to ^~^40°C in a benchtop hot block.

For rat tail collagen samples molten LMP agarose (^~^40 μL) was placed onto the pedestal of an imaging chamber, and pieces of rat tail were placed inside the agarose bead using fine-tip tweezers. The imaging chamber was then placed in a fridge at 4°C for 5 minutes to gel.

Cell Spheroid samples were removed from storage media and placed in a 1.5 mL Eppendorf with ^~^80 μL LMP agarose. 40μL of agarose containing the sample was removed by pipette, placed on the PDMS pedestal, and was inspected to check that the sample was approximately central in the agarose bead. The imaging chamber was then inverted to ensure the spheroid would settle near the surface of the agarose, and the chamber was placed in a fridge at 4°C for 5 minutes to gel.

For oocytes, the samples were inserted with a glass capillary into a droplet of 37°C 0.5% LMP agarose prepared freshly with M2 media just prior to the agarose gelling.

In all cases the imaging chamber was then placed on the microscope translation stage and was covered with water or media prior to imaging.

### Sample Staining

Stock solutions of FM4-64 (3 mM, Insight Bio, CAS No: 162112-35-8) were prepared by dissolution in DI water. A working solution was prepared by 1:300 dilution in M2 media to yield a final concentration of ^~^10 μM. Oocytes were incubated in the dye for 10 minutes before imaging. Hoechst 33342 (Sigma-Aldrich) was prepared as a 20mg/mL stock in H_2_O and diluted 1:10,000 into the imaging media prior to use.

### Beam Profile Simulations

Beams profiles were simulated using a Matlab script adapted from the Optical Modelling Group at St Andrew’s^63^. Scripts were updated to account for wavelength-dependence of the input phase mask α-value and rotation of the input phase profile about the optical axis. One- and two-photon fluorescence input beam profiles were simulated for selected angles of the input phase mask.

### Beam Profile Measurements

A dilute solution of FITC (10 μM) in water was used for visualisation, alignment and measurement of focussed beam profiles. The useful width of each beam profile was estimated using the line profiler in ImageJ. We used 1/e^2^ of the maximal value to define the profile limits. For 1PF beam profile measurement, λ_ex_ = 488 nm, λ_em_ = 520 ± 20 nm, P_ex_ = <1 μW, t = 100 ms. For 2PF beam profile measurement, λ_ex_ = 720 nm, λ_em_ = 520 ± 20 nm, P_ex_ = <1 mW, t = 100 ms.

### Resolution Measurements

Fluorescent microbeads (0.1 μm, FluoSpheres, F8803) were diluted 2000x and suspended in 1% agarose. Using either 488 nm CW laser excitation or 860 nm fs-pulsed laser excitation for 1PF or 2PF respectively, 3D image stacks were acquired to a depth of 40 μm with a slice separation of 100 nm. Image stacks were process using custom-written Matlab scripts. In brief, images were imported to the workspace, and local maxima were found. Maxima above a given intensity threshold removed background noise, and gave the positions of the fluorescent beads. Intensity profiles were taken across 30 px in x,y,z centred on the fluorescent bead maximum for all bead positions. A Gaussian curve was fitted to each intensity profile, and FWHM values were extracted from this fitting. Measures of the mean and standard deviation of FWHM values were taken in x and y, and separately in z, to find an overall measure of the lateral and axial resolution of the microscope.

### Wavelength Scans

Wavelength scans were performed by automated control of the fs-pulsed laser (MaiTai-BB, SpectraPhysics) from within our custom microscope control GUI (Python 2.7). Images were acquired for illumination wavelengths in the specified range (typically 730 to 870 nm).

### K-Means Cluster Analysis

Wavelength-scan image stack (x,y,λ) was imported into the Matlab workspace with 2×2 binning in xy, and the minimum image value was subtracted from each image to remove the image background offset. After reorganisation into a 2D matrix (p,λ) where p indicates the x,y position, K-means clustering into 8 clusters was performed on the data using the in-built Matlab function. Average spectra from each cluster were normalised, and clusters were manually grouped based on their spectral similarity. The cluster vectors were reorganised into false-colour images, which indicated the spectrally distinct nature of different regions within the sample.

### Image Processing

Images were processed using FIJI^64^, using standard packages. Where contrasts were adjusted all images in one panel were treated in the same way.

### Statistical Tests

Student’s t-test and ANOVA statistical tests were performed on measurements from SHG images (**Fig1 f and g**). Tests were performed in Graphpad Prism (8.2.1), and P < 0.05 were considered to be statistically significant. All figures show 5-95% limits.

### Figure Preparation

Graphs were prepared using Prism Graphpad, micrographs were prepared using FIJI^64^, and overall figure assembly was performed in Adobe Illustrator.

## Supporting information

SHG-LSM Supplementary Figures and Table

## Acknowledgements

We thank M2 Life for provision of the Light Sheet Microscope, and Neveen Hosny, Gunnar Spickermann, Robert Forster and Graeme Malcolm at M2 Life for their input and discussions.

## Funding

NH acknowledges funding by EPSRC Case Conversion studentship (EP/N509747/1) co-funded by M Squared. PJ is co-funded by EPSRC Doctoral Training grant (EP/N509747/1) and ERC grant NanoChemBioVision (638258). SL is funded by Wessex Medical Research (Z08) and EPSRC Impact Acceleration Account, University of Southampton. MGJ acknowledges the British Lung Foundation (SRG19\100001). SM acknowledges the European Research Council (ERC) grant NanoChemBioVision (638258) and EPSRC grant (EP/T020997/1).

## Author Contributions

NH, SL and SM led the design of experiments and analysis. LSM was designed and built by NAH and GS with input from GM and RF. Modification of LSM system for SHG and characterisation was performed by NH and SL. Murine oocytes were provided by SL, and rat tail collagen was provided by PJ. CB, RR, ED and MJ provided lung fibroblast spheroids. NH performed SHG-LSM imaging experiments. Data and image processing was performed by NH and SL with input from SM, and from CB, and MJ on LF spheroid data. Beam profile simulations performed by NH with input from KB. NH, SL and SM wrote the first draft of the manuscript with inputs from all authors. NH, SL and SM led the finalisation of the manuscript. All authors contributed to and approved the final version.

## Corresponding author

Prof Sumeet Mahajan

## Ethics declarations

### Competing interests

The authors declare no competing interests.

## Supplementary information

List of supplementary information/figures:

Supplementary Figure 1: Simulations of Rotated Airy Beam

Supplementary Figure 2: Experimental measurement of LSM FOV and resolution using 2PF

Supplementary Figure 3: SHG from Rat Tail Collagen with changing input polarisation state

Supplementary Figure 4: SHG from Rat Tail Collagen with circular polarised illumination

Supplementary Video 1: Label-free Multimodal imaging of Lung Fibroblast Spheroid

## Notes

### Competing Interest Statement

The authors have declared no competing interest.

