## Supplementary material for "Label-free and Multimodal Second Harmonic Generation Light Sheet Microscopy": SHG-LSM Supplementary Figures and Table

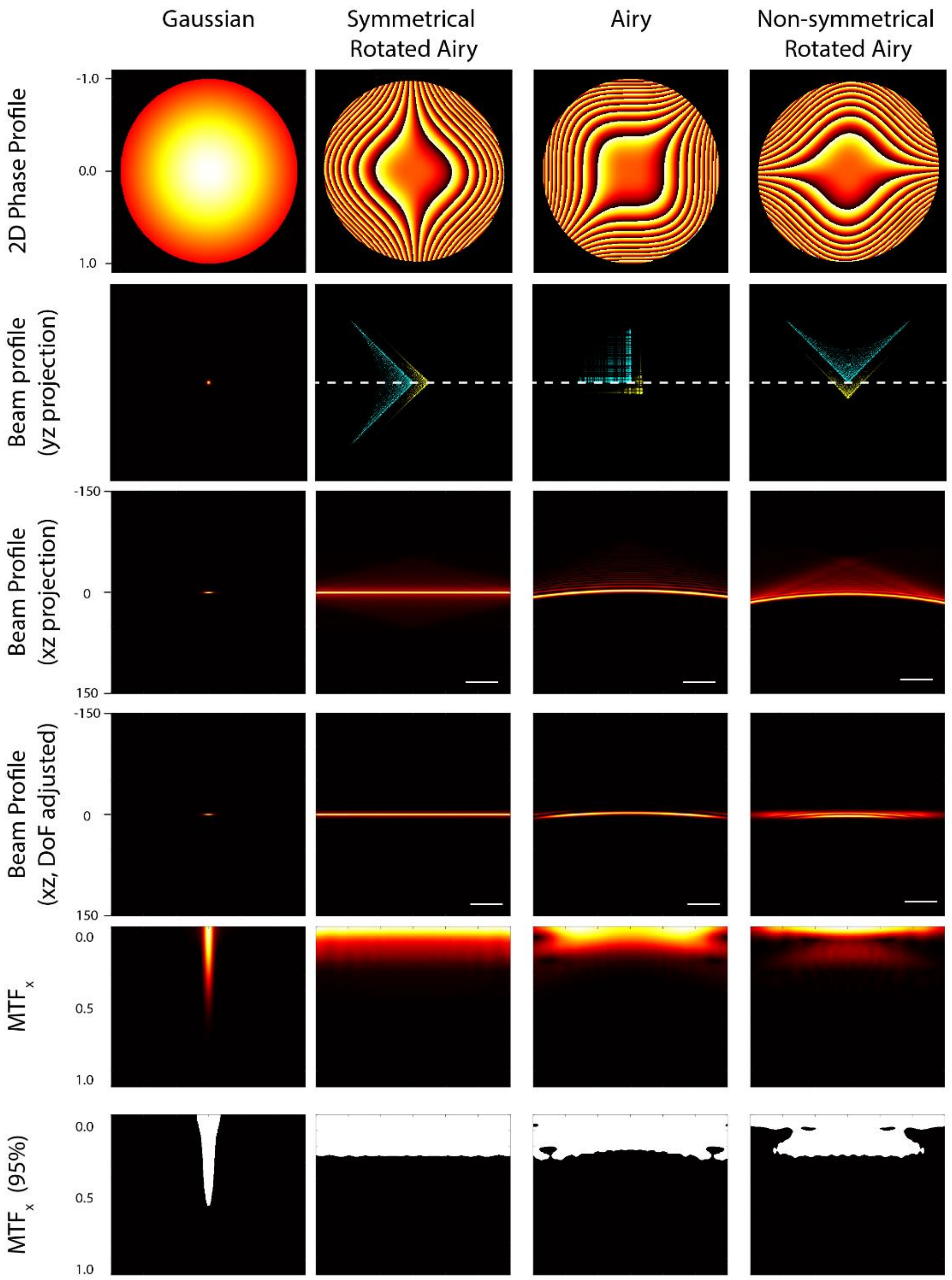


**Figure S1** ***Beam profile simulations for Rotated Airy Light Sheet Microscopy*** *Simulations were performed for Gaussian (G), Symmetrical Rotated Airy (SRA), Airy (A), and Non-Symmetrical Rotated Airy (NSRA) beam profiles. (a) The 2D phase profiles projected onto the objective back aperture, wrapped between* $-\pi^C$ *and* $\pi^C$*. (b) Focussed beams profile in yz from the 2D Fourier transform of the 2D phase profiles, showing rotation of the Airy lobes about the propagation axis, and (c) beam profile projection in xz showing extent of beam across the 300 µm FOV. The main lobe moves out of imaging plane for NSRA and A, only for SRA does it travel within the imaging plane. (Cyan – beam cross section at centre of focus, Yellow – 50 µm from centre of focus). (d) A finite region of the NSRA and A beam profile main lobe remains within the DoF of the objective, though this depends on chosen α-value. The main lobe of the SRA beam profile remains in the DoF of the detection objective across the FOV. (e,f) Modulation Transfer Function and MTF 95% contrast level. This is the 2D Fourier transform of the PSF along the x axis, indicating the achievable contrast at each position in x. Though the highest contrast can be achieved using the high-NA Gaussian beam profile, this is over a very small range in z. The MTF for the SRA beam is approximately uniform, providing the best high-resolution imaging capability across the widest FOV.*

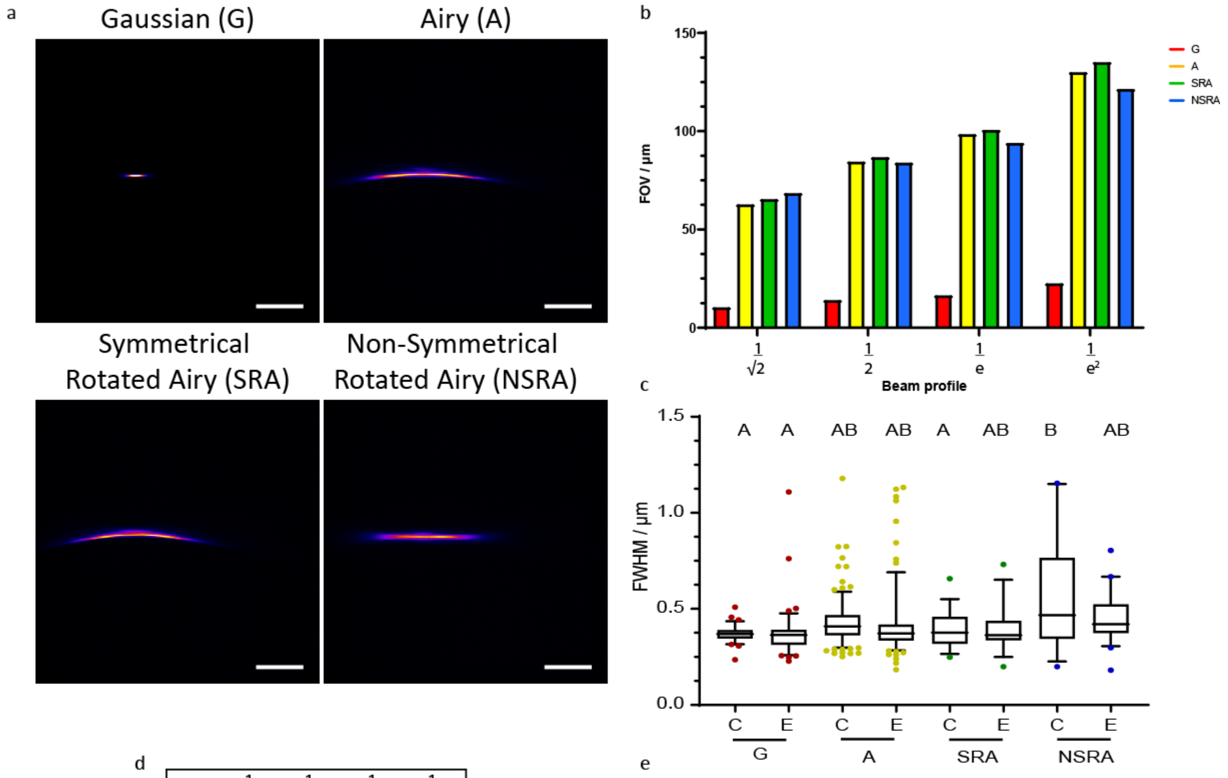


d

| | $\frac{1}{\sqrt{2}}$ | $\frac{1}{2}$ | $\frac{1}{e}$ | $\frac{1}{e^2}$ |
|---|---|---|---|---|
| G | 10.3 | 62.7 | 65.4 | 68.5 |
| A | 14.0 | 84.5 | 86.7 | 83.9 |
| RA1 | 16.4 | 98.4 | 100.6 | 94.0 |
| RA2 | 22.6 | 130.0 | 135.1 | 121.4 |

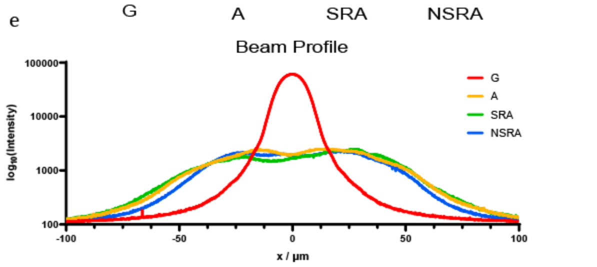


***Figure S2 Experimental two-photon fluorescence beam profiles and resolution*** *(a) Fluorescence emission from fluorescein in the FOV of the microscope excited using fs-pulsed laser with Gaussian, Airy, Rotated Airy 1 and Rotated Airy 2 beam profiles. (b) Comparing useable FOV of each beam type with different metrics. Using FWHM,* $\frac{1}{e}$ *or* $\frac{1}{e^2}$ *measurement of useable FOV, SRA (used in this work) gives the largest useable FOV, and all Airy-type beams give an approx. 6.5x increase in the effective useable FOV compared to the Gaussian beam. (c) FWHM for beads in 4 regions from* $1/e^2$ *to 0 either side of beam centre. A significant difference in FWHM for NSRA was measured for fluorescent beads in the 50 µm range about the beam focus using one-way ANOVA, no significant difference in lateral resolution for remaining beam types. (d) Data table of values in (b). (e) Intensity profiles along the beam axis (G, NSRA), or along the main lobe (A, SRA).*

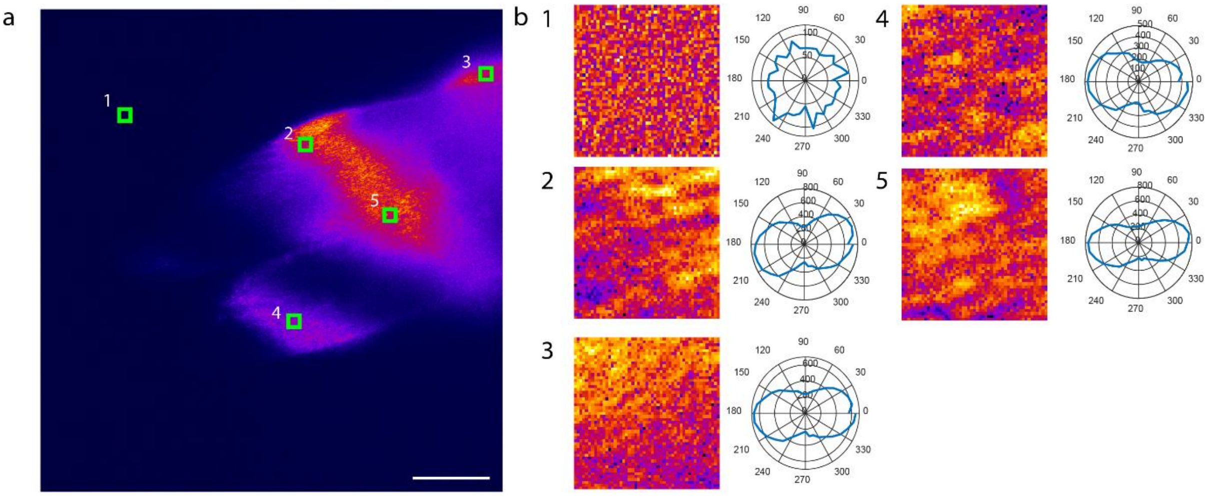


***Figure S3 SHG signal from Rat Tail Collagen changes with rotation of input polarisation*** *(a) Rat tail tendon orientated approximately along the propagation axis was imaged with polarisation-resolved SHG microscopy (Image shows signal with polarisation fast axis in the plane of the light sheet). Fibres orientated in some direction pointing away from the detection objective do not produce strong SHG signal for any polarisation (see dark region between 4 and 5) (b) Differences in SHG signal with changing polarisation angle can be observed by extracting small regions from the main image. The small differences in polarisation angle relate to the 3D orientation of the collagen fibres. Collagen signal is moderately anisotropic.*

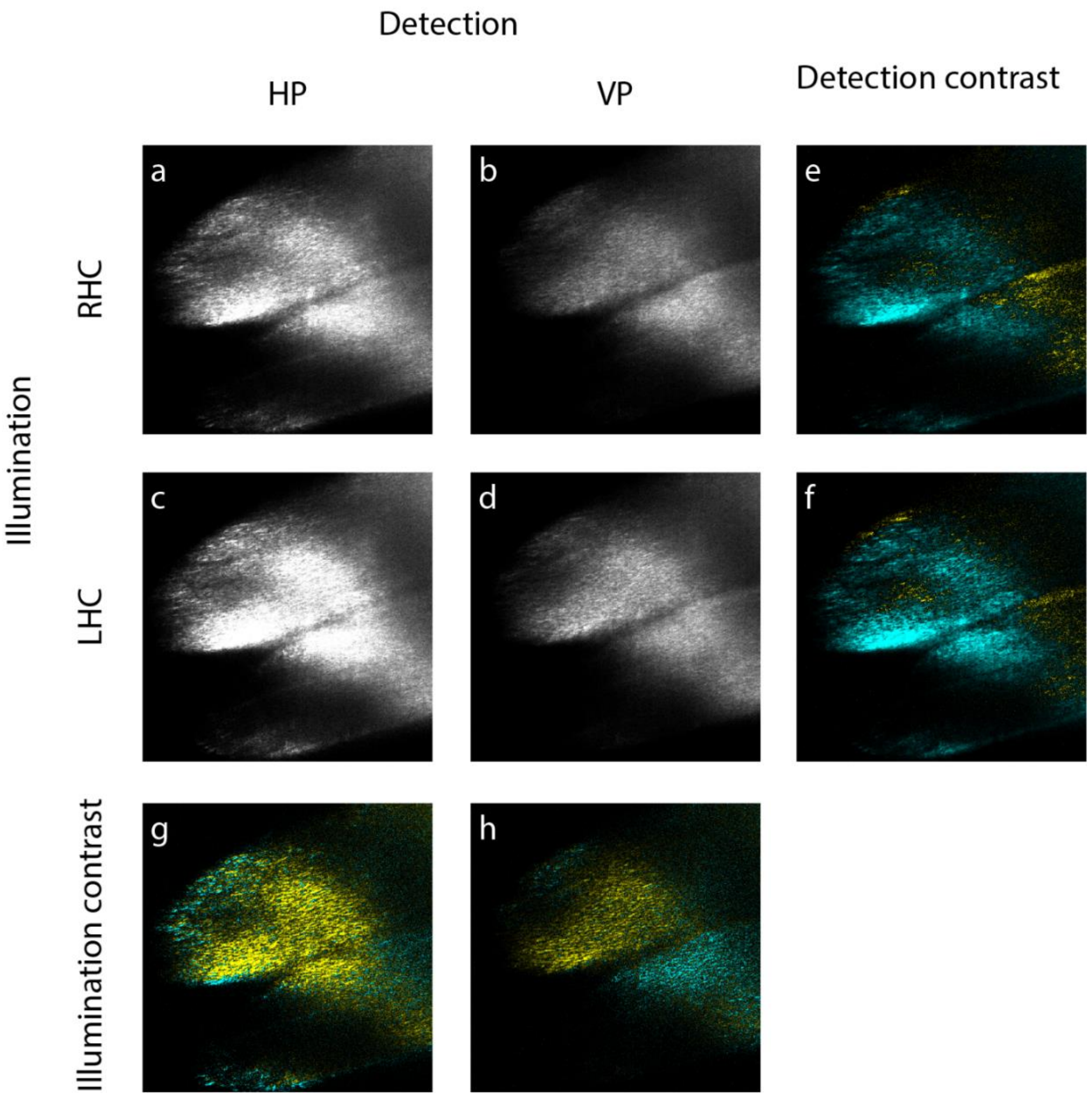


***Figure S4. SHG Circular polarisation contrast*** *Spatial differences in SHG signal with LHC and RHC polarisation in illumination path. Further differences are observed using linear polarisation contrast in detection path. Images in Detection contrast created using thresholded HP-VP (cyan) and VP-HP (yellow) images. Images in Illumination contrast created using LHC-RHC (yellow) and RHC-LHC (Cyan).*

*Video S1: Label-free Multimodal Imaging of Lung Fibroblast Spheroid (SHG/2PF)*

**Supplementary Video 1: Label-free Multimodal imaging of Lung Fibroblast Spheroid** Front section of a lung fibroblast spheroid was imaged to a depth of 140 µm across a 180µm x 300µm effective FOV. Two-photon excited cellular autofluorescence (red) and SHG from collagen in ECM (cyan) provide complementary signals that are spectrally distinct and readily combined with one-photon fluorescence and other imaging modalities. SHG-LSM can be used for imaging highly scattering biological samples up to a depth of >140 µm.

*Supplementary Table 1: Imaging conditions for each experiment*

| Experiment | 1 – Beam profile measurements | 2 – Rat tail collagen SHG polarisation contrast | 3 – SHG wavelength scan |
|---|---|---|---|
| Figures | 1, S2 | 2, S4 | 2 |
| Imaging mode(s) | 2PF | SHG | SHG, 2PF |
| Objectives | (Illumination) Olympus UMPLFLN20XW, 20x, NA 0.5, WD 3.5;<br>(Detection) Olympus LUMPLFLN40XW, 40x, NA 0.8, WD 3.3 | | |
| Excitation lasers | Spectra-Physics MaiTai-BB 710-990 nm, 100 fs, 80MHz | | |
| Excitation power and wavelength | 18 µW (488nm) ; <5 mW (MaiTai) | 190 mW (740 nm), 272 mw (800 nm) | Between ~160 mW (710 nm) and 280 mW (820 nm) |
| Detection path optical filters / nm | 520 ± 20 nm (2PF) ; 400 ± 20 nm (SHG) | | 405 ± 10 nm (SHG) |
| Illumination path polarisation elements | None | Achromatic half-wave plate / Quarter-wave plate | Achromatic half-wave plate |
| Polarisation state (in) | - | Parallel / Perpendicular / Circular | Parallel / Perpendicular |
| Detection path polarisation elements | None | None, Linear polariser | None, Linear polariser |
| Camera | Hamamatsu Orca Flash 4.0 V2.0 sCMOS (2048 x 2048 px) | | |
| Imaging Mode | Water cooled (20°C), MAX, Global readout | | |
| Camera exposure / ms | 100 | 200, 500 | 200, 500 |
| Sample condition | FITC (<1µM) in DI water solution | PFA-fixed rat tail collagen in 0.5% low-melting point agarose | |
| Post-processing steps | Despeckle, background subtraction | | |

| Experiment | 4 – SHG polarisation scan with Rat Tail Collagen | 5 – Oocyte autofluorescence and SHG membrane dye | 6 – Lung fibroblast spheroids |
|---|---|---|---|
| Figures | S3 | 3, 4 | 5 |
| Imaging mode(s) | SHG | 2PF, SHG, 1PF | SHG, 2PF |
| Objectives | (Illumination) Olympus UMPLFLN20XW, 20x, NA 0.5, WD 3.5;<br>(Detection) Olympus LUMPLFLN40XW, 40x, NA 0.8, WD 3.3 | | |
| Excitation lasers | Spectra-Physics MaiTai-BB 710-990 nm, 100 fs, 80MHz | Spectra-Physics MaiTai-BB 710-990 nm, 100 fs, 80MHz;<br>488 nm CW laser (up to 15 mW) | Spectra-Physics MaiTai-BB 710-990 nm, 100 fs, 80MHz |
| Excitation power and wavelength | Between ~190 mW (710 nm) and 272 mW (820 nm) | 190 mW (740 nm), 272 mW (800 nm) | Between ~280 mW (710 nm) and 400 mW (820 nm) |
| Detection path optical filters / nm | 520 ± 20 nm (2PF) ; 405 ± 10 nm (SHG) | 520 ± 20 nm (2PF) ; 405 ± 10 nm (SHG); 710 ± 10 nm (1PF) | 520 ± 20 nm (2PF) ; 405 ± 10 nm (SHG) |
| Illumination path polarisation elements | Achromatic half-wave plate | | |
| Polarisation state (in) | Parallel / Perpendicular | Parallel / Perpendicular | Perpendicular |
| Detection path polarisation elements | None / Linear polariser | None / Linear polariser | None |
| Camera | Hamamatsu Orca Flash 4.0 V2.0 sCMOS (2048 x 2048 px) | | |
| Imaging Mode | Water cooled (20°C), MAX, Global readout | | |
| Camera exposure / ms | 20-200 | 1000, 200 | 1000 (2PF), 4000 (SHG) |
| Sample condition | PFA-fixed rat tail collagen in 0.5% low-melting point agarose | Murine oocytes with FM4-64 dye (10 µM) bound to membrane after treatment, in 0.5% low-melting point agarose | 3D cell culture of PFA-fixed lung fibroblasts in 0.5% low-melting point agarose |
| Post-processing steps | Despeckle, background subtraction | Despeckle, background subtraction, K-Means clustering | Despeckle, background subtraction |
